# An RNA-guided genome editing tool for mammalian cells and embryos: Universal CRNA-guided GE AssemblY (UCAY) system

**DOI:** 10.64898/2026.11.26.769381

**Authors:** Suong Thi Nguyen, So Makabe, Hideyuki Kodama, Yuichi Kobashi, Yumu Yaku, Yuichiro Nakayama, Mao Matsushita, Aya Nakai, Maki Hirata, Megumi Nagahara, Ikuo Nakamura, Yasuhiro Moriizumi, Fuminori Tanihara

## Abstract

Genome editing technologies, such as CRISPR/Cas9, have revolutionized life science research. However, their application can be constrained by protospacer-adjacent motif (PAM) requirements and complex intellectual property frameworks. Here, we established an RNA-guided genome editing platform, Universal CRNA-guided GE AssemblY (UCAY), which uses a FokI–Nova fusion protein recruited to target sequences through guide RNA containing UCAY motifs. We evaluated the UCAY-mediated cleavage activity in a PCR-amplified fragment containing the target site and mammalian cell lines using GFP-expressing HEK293 cells. Introduction of GFP-targeted UCAY components by electroporation disrupted GFP fluorescence, suggesting genome-editing activity in cultured cells. We examined whether UCAY could be applied to mammalian embryos using in vitro-fertilized porcine zygotes targeting the *GGTA1* gene. Cytoplasmic microinjection of UCAY ribonucleoprotein complexes induced mutations in up to 73.9% of blastocysts without impairing embryonic development. These findings demonstrate the feasibility of UCAY-mediated genome editing in mammalian cells and embryos and establish UCAY as a versatile alternative platform for future biomedical and industrial applications.

## INTRODUCTION

Genome editing technologies have rapidly advanced over the past two decades, enabling the precise modification of nucleic acid sequences in many organisms. Early programmable nucleases included zinc finger nucleases (ZFNs) [1, 2], which consisted of engineered zinc finger DNA-binding domains fused to a nonspecific DNA cleavage domain, and transcription activator-like effector nucleases (TALENs) [3], which combined DNA-binding modules derived from *Xanthomonas* transcription activator-like effectors with a nuclease domain. These systems show the feasibility of sequence-specific genome modification but require complex protein engineering for each target site.

The emergence of the CRISPR/Cas9 system has considerably improved the efficiency and accessibility of genome editing [4, 5]. This system is composed of a guide RNA (gRNA), which contains a sequence complementary to the target DNA, and Cas9 protein, which forms a complex with the gRNA to induce site-specific DNA cleavage. Because target recognition relies on RNA–DNA base pairing, CRISPR/Cas9 is substantially easier to design and implement than ZFNs and TALENs. Despite this, CRISPR/Cas systems have inherent limitations. One major constraint is the requirement for a protospacer-adjacent motif (PAM) sequence near the target site, which restricts the range of editable loci within the genome. The licensing complexities associated with CRISPR technologies may limit their broad application [6]. Therefore, there is a strong need for alternative genome editing platforms that do not depend on PAM sequences and allow for the flexible targeting of nucleic acids.

Here, we developed a novel genome editing platform termed “Universal CRNA-guided GE AssemblY (UCAY),” based on a design concept in which RNA specifies the target location while a separate effector protein executes the desired function. Unlike CRISPR/Cas systems, UCAY functions as a modular platform that recruits arbitrary functional proteins to define nucleic acid sequences under RNA guidance. The UCAY system comprises two main components. The guide RNA contains (i) hybridization regions that bind to sequences flanking the target site, and (ii) a guide region that recruits a fusion protein. The fusion protein comprises an RNA-binding domain that recognizes the guide region and an effector domain that is responsible for modifying the target nucleic acid.

As a proof of concept, we engineered a fusion protein by combining the FokI nuclease domain with the RNA-binding protein Nova-1 (FokI–Nova fusion protein) [7]. Nova-1 is a neuron-specific nuclear RNA-binding protein, and its KH3 domain specifically recognizes three repeats of a four-nucleotide sequence (U, C, A, and Y; where Y denotes a pyrimidine, U or C) within the pre-mRNA of the α2 glycine receptor (α2-glyR). Based on this property, we designed a UCAY guide RNA (UgRNA) by linking a sequence complementary to the target gene to two UCAY repeat motifs derived from α2-glyR. We propose that one FokI–Nova fusion protein, connected via a linker sequence, binds to each UCAY repeat sequence through RNA–protein interactions. This arrangement is expected to bring the two FokI nuclease domains into proximity, allowing dimerization and double-stranded DNA cleavage in a manner analogous to that of ZFNs and TALENs. Using this system, we showed that the combination of UgRNA and FokI–Nova enables the modification of target nucleic acid sequences in mammalian cells and fertilized embryos without dependence on PAM sequences. These findings established UCAY as a versatile and programmable platform for nucleic acid editing.

## RESULTS

### Experiment 1: Construction and in vitro validation of the UCAY system

A schematic representation of the UCAY system using the FokI–Nova fusion protein is shown in Figure 1A. The complete nucleotide sequence of the FokI–Nova expression plasmid, plasmids used for UgRNA transcription, and amino acid sequence of the FokI–Nova fusion protein are provided in Supplementary Data S1 and S2, respectively. UgRNA targeting the GFP sequence (UgRNA-GFP) was designed to evaluate the cleavage activity of the UCAY system in vitro (Supplementary Figure S1). UgRNA-GFP (200 ng) was incubated with the PCR-amplified GFP fragment (714 bp) containing the target site (Supplementary Figure S2) (200 ng) and FokI–Nova protein (125 ng). Following agarose gel electrophoresis, DNA fragments likely representing cleavage products were extracted from the gel. The recovered DNA fragments were cloned using a Quick Blunting Kit (New England Biolabs, Ipswich, MA, USA), and 102 clones were sequenced. Sequence analysis identified two major cleavage sites that were generally consistent with the banding patterns observed by analysis using the microchip electrophoresis system (MultiNA) (Figure 1B, C, D), indicating that the hybridization region of UgRNA successfully bound to the GFP target sequence and guided site-specific cleavage. These results indicate that the UCAY system, mediated by the FokI–Nova fusion protein, can cleave target nucleic acid sequences independently of PAM constraints.

**Figure 1.**
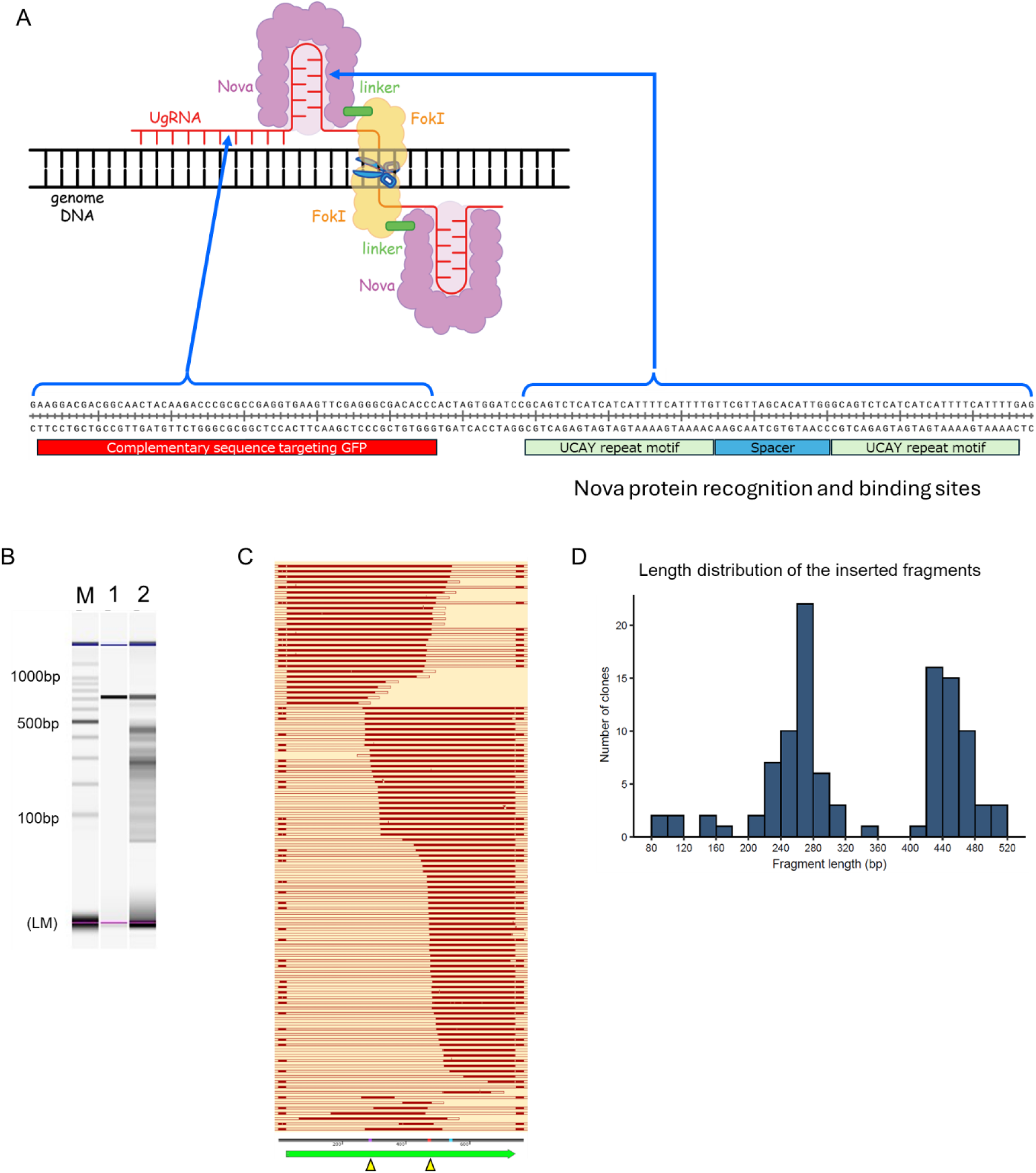
Overview of the UCAY system and validation of its target sequence cleavage activity. (A) A schematic representation of the UCAY system using the FokI–Nova fusion protein. (B) The cleavage activity of UgRNA targeting the GFP sequence was analyzed using MultiNA. The lanes in the electrophoresis image are as follows; M: 100 bp ladder. Lane 1: GFP PCR product only. Lane 2: GFP PCR product + FokI–Nova fusion protein + UgRNA. (C) Sequence analysis of individual clones visualized using SnapGene software. To allow alignment regardless of the orientation of the cloned inserts within the vector, a .dna file containing only the GFP region and its flanking sequences was generated, and the sequencing results from individual clones were aligned using the SnapGene alignment function. The full-length target GFP amplicon (714 bp) is shown as the light-green arrow at the bottom, whereas the inserted fragments corresponding to UCAY-mediated cleavage products are indicated by dark-brown lines. The major putative cleavage sites are indicated by yellow triangles. (D) Histogram showing the length distribution of the inserted fragments.

### Experiment 2: Genome editing in mammalian cell lines using UCAY

We evaluated whether the UCAY system can induce genome editing in mammalian cells. Human embryonic kidney (HEK) 293 cells stably expressing GFP (293/GFP, AKR-200; Cell Biolabs, CA, USA) were used as a model system. UgRNA-GFP (100 ng/µl) and FokI–Nova protein (50 ng/µl) were introduced into cells via electroporation using the CUY21EDIT II system (BEX, Tokyo, Japan). After 4 d, fluorescence microscopy showed a loss of the GFP signal in a subset of cells (Figure 2A), suggesting successful disruption of the GFP gene. Fluorescence-activated cell sorting (FACS) analysis indicated that 1.27% of the cells lost GFP fluorescence, showing that the genome editing activity was mediated by the UCAY system (Figure 2B, C). Although direct comparison is difficult because the electroporation conditions and the amounts or concentrations of the individual components differed among studies, the editing efficiency observed here was lower than that typically reported for CRISPR/Cas9 ribonucleoprotein (RNP)-based systems in human cells [8, 9]. In both these studies, Cas9 protein and gRNA were premixed to form RNP complexes before being introduced into cells by electroporation. Nevertheless, the present results confirmed that UCAY-mediated cleavage is functional in mammalian cell lines.

**Figure 2.**
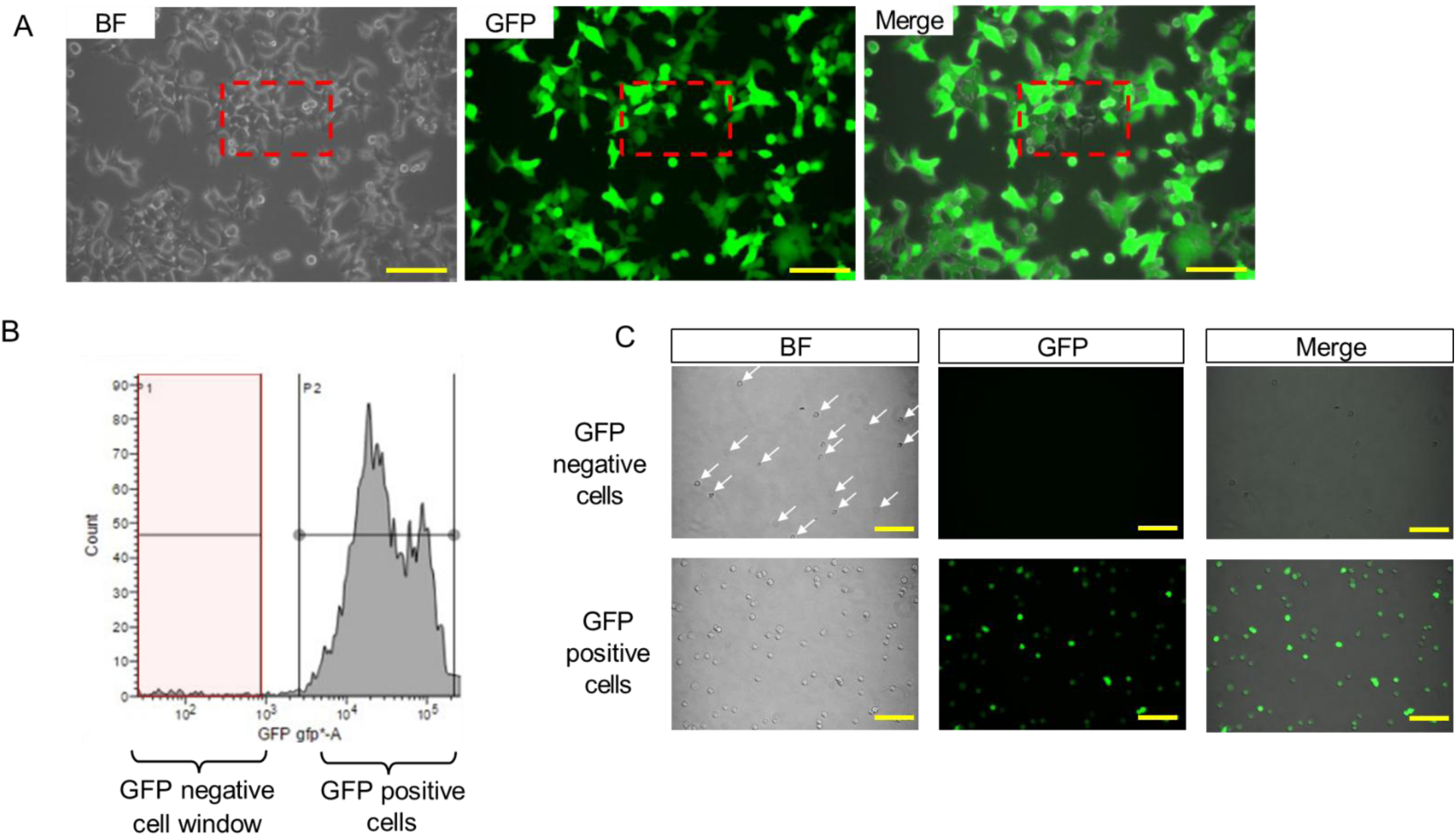
Genome editing in cell lines using the UCAY system composed of the FokI–Nova protein and a UgRNA-GFP. (A) Genome editing of cell lines constitutively expressing GFP introduced GFP-targeting UgRNA and FokI–Nova proteins via electroporation. Regions enclosed by red squares indicate colonies in which GFP fluorescence was lost. The scale bar in each panel represents 100 µm. (B) FACS analysis of GFP-negative and GFP-positive cells after UCAY-mediated genome editing targeting GFP. (C) Fluorescence images of putative GFP-negative and GFP-positive cells separated by FACS. The scale bar in each panel represents 100 µm.

### Experiment 3: Genome editing in porcine zygotes using UCAY

Genetically modified pigs are ideal donors for pig-to-human xenotransplantation [10–12]. Traditionally, genetically modified pigs are generated via somatic cell nuclear transfer using modified donor cells, which is a technically demanding and complex process. The advent of genome-editing technologies has enabled the direct modification of zygotes, simplifying the production of genetically engineered pigs. Given the shortage of human donor organs, pigs have become increasingly recognized as promising alternatives. However, the presence of xenoantigens remains a major barrier to xenotransplantation. One of the primary xenoantigens is the galactosyl-α1,3-galactose (Galα(1,3)Gal) epitope, synthesized by α1,3-galactosyltransferase encoded by the *GGTA1* gene [12, 13]. Therefore, we designed a UCAY system targeting *GGTA1*.

Two UgRNAs (UgRNA1 and 2) were designed to target *GGTA1* exon 2 (Figure 3A and Supplementary Figure S1). Each UgRNA (100 ng/µl) was co-injected with FokI–Nova protein (100 ng/µl) into in vitro-fertilized porcine zygotes via cytoplasmic microinjection. The injection concentrations were determined based on a previous CRISPR/Cas9 study [14]. Control groups included non-injected embryos, solvent-injected embryos, and embryos injected with CRISPR/Cas9 RNP complexes (100 ng/μl gRNA and 100 ng/μl Cas9 protein) targeting *GGTA1* exon 2 (Figure 3A).

**Figure 3.**
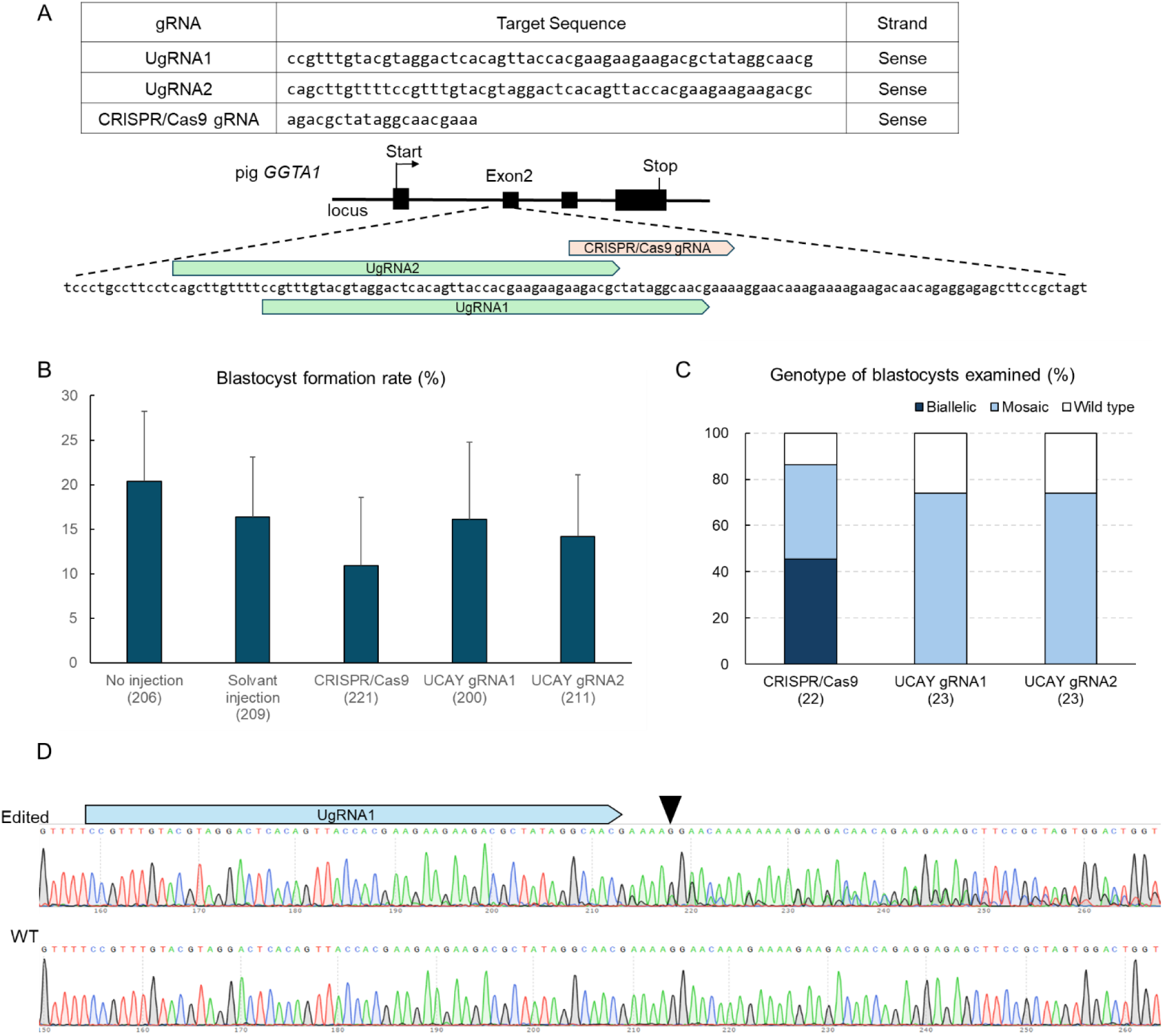
Genome editing via cytoplasmic microinjection of the UCAY system composed of the FokI–Nova protein and a UgRNA targeting GGTA1. (A) UgRNA target sequences and the genomic structure of the *GGTA1* locus. The target sequence of the CRISPR/Cas9 gRNA used as control is also shown. (B) Rate of blastocyst formation in microinjected zygotes. For each treatment group, four replicates of 200–221 oocytes per treatment were analyzed. Numbers in parentheses indicate the total number of treated zygotes. Values of means ± SEM are shown. (C) Genotypes of blastocysts after cytoplasmic microinjection with FokI–Nova protein and each UgRNA targeting *GGTA1* or CRISPR/Cas9 RNP targeting *GGTA1*. The percentage of blastocysts was defined as the ratio of the mutant blastocysts to the total number of blastocysts. The percentages of mutant blastocysts were analyzed using the chi-square test. WT, wild type; Biallelic, biallelic mutant; Mosaic, mosaic mutant. Numbers in parentheses indicate the total number of blastocysts analyzed. (D) Representative genomic sequences of blastocysts formed after cytoplasmic microinjection of FokI–Nova protein and UgRNA1. Arrowheads indicate the putative UCAY cleavage sites.

No significant differences in blastocyst formation rates were observed among the injection groups (Figure 3B), indicating that the UCAY components did not impair early embryonic development. Blastocyst genotyping was performed using Sanger sequencing, followed by TIDE analysis [15]. Blastocysts in which no wild-type (WT) alleles were detected were classified as biallelic mutants, whereas those containing both WT and mutant alleles were classified as mosaic. All embryos edited using UgRNA1 and UgRNA2 exhibited mosaicism. However, mutations were detected in 73.9% of blastocysts for both UgRNAs (Figure 3C). Representative sequencing results are shown in Figure 3D.

We also evaluated the plasmid-based delivery in zygotes. All-in-one plasmids encoding UgRNA and FokI–Nova (pUgRNA1-GGTA1-UCAY and pUgRNA2-GGTA1-UCAY) (Supplementary Data S1F, G) were microinjected into zygotes at a concentration of 10 ng/µl. As a control, the group was injected with the solvent (distilled water). Similar to the RNP injection, no significant differences in blastocyst formation rates were observed in the injection group (Figure 4A). Using UgRNA1 and UgRNA2, all embryos remained mosaic with mutation rates of 16.7% and 11.5%, respectively (Figure 4B).

**Figure 4.**
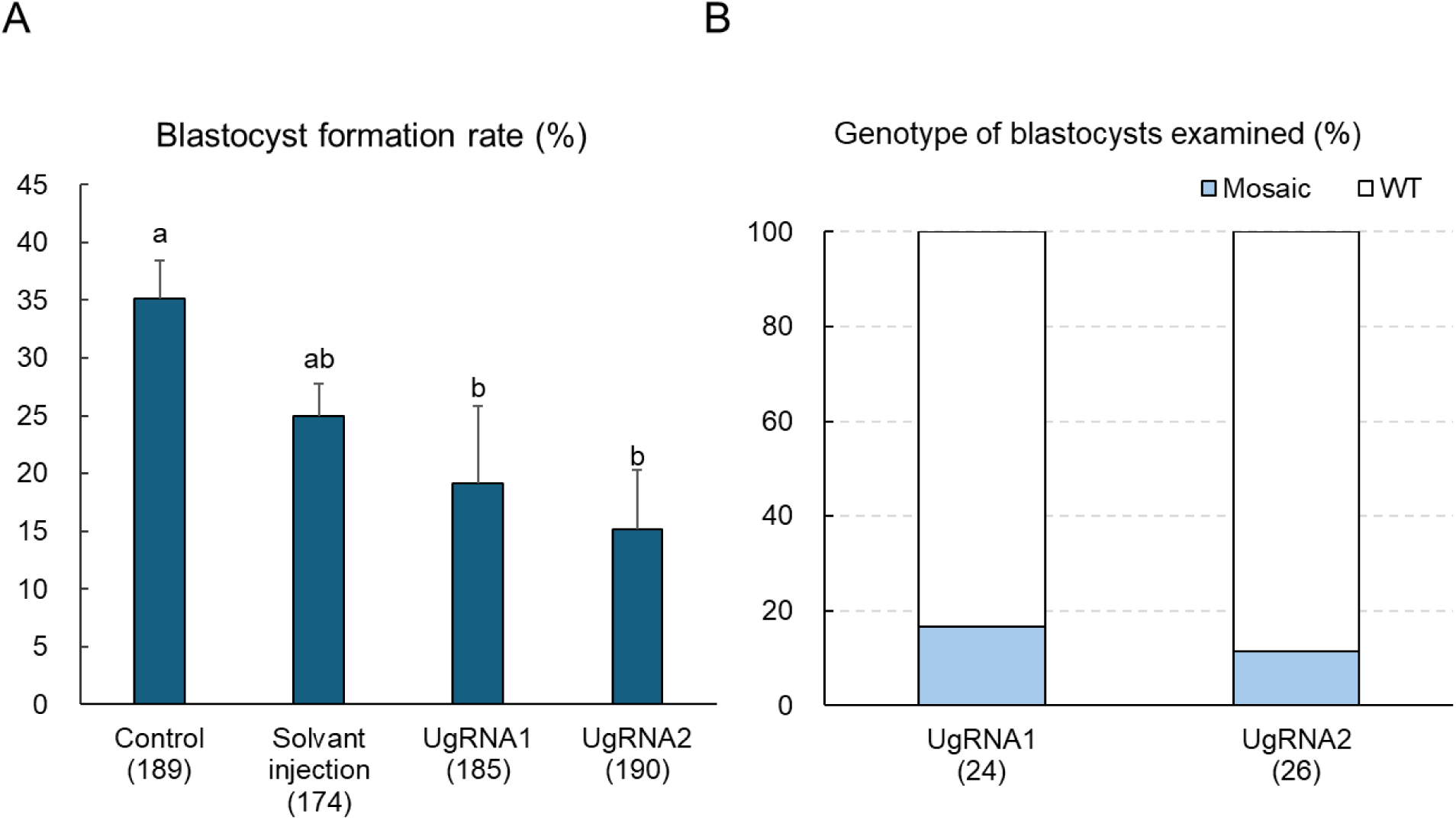
Genome editing via cytoplasmic microinjection of an all-in-one plasmid expressing FokI–Nova and UgRNA targeting GGTA1. (A) Rate of blastocyst formation in microinjected zygotes. For each treatment group, four replicates of 174–190 oocytes per treatment were analyzed. Numbers in parentheses indicate the total number of treated zygotes. ^a–b^ Values with different superscripts differ significantly (*p* < 0.05). Values of means ± SEM are shown. (B) Blastocyst genotypes after cytoplasmic microinjection of FokI–Nova and UgRNA targeting *GGTA1* expression all-in-one plasmid. The percentage of blastocysts was defined as the ratio of the mutant blastocysts to the total number of blastocysts. The percentages of mutant blastocysts were analyzed using the chi-square test. WT, wild type; Mosaic, mosaic mutant. Numbers in parentheses indicate the total number of blastocysts analyzed.

Compared to cell line experiments, higher genome editing efficiencies were observed in zygotes. This difference may be attributed to using cytoplasmic microinjection rather than electroporation. However, the underlying mechanism remains unclear and warrants further investigation. To further evaluate the delivery methods, we introduced UgRNA and the FokI–Nova protein into porcine zygotes via electroporation. Electroporation was performed as previously reported [16]. Each UgRNA targeting *GGTA1* (100 ng/µl) and FokI–Nova protein (100 ng/µl) were introduced into in vitro-fertilized porcine zygotes by electroporation.

As with microinjection, no significant differences in blastocyst formation rates were detected among the experimental groups (Figure 5A), indicating that electroporation-based delivery did not adversely affect early embryonic development. Compared to the microinjection groups, the blastocyst formation rate was maintained at a higher level, indicating that the invasiveness of the electroporation procedure into the embryos was extremely low. Genotyping analysis showed that all embryos exhibited mosaic mutations, with mutation rates of 23.3% and 9.7% (Figure 5B).

**Figure 5.**
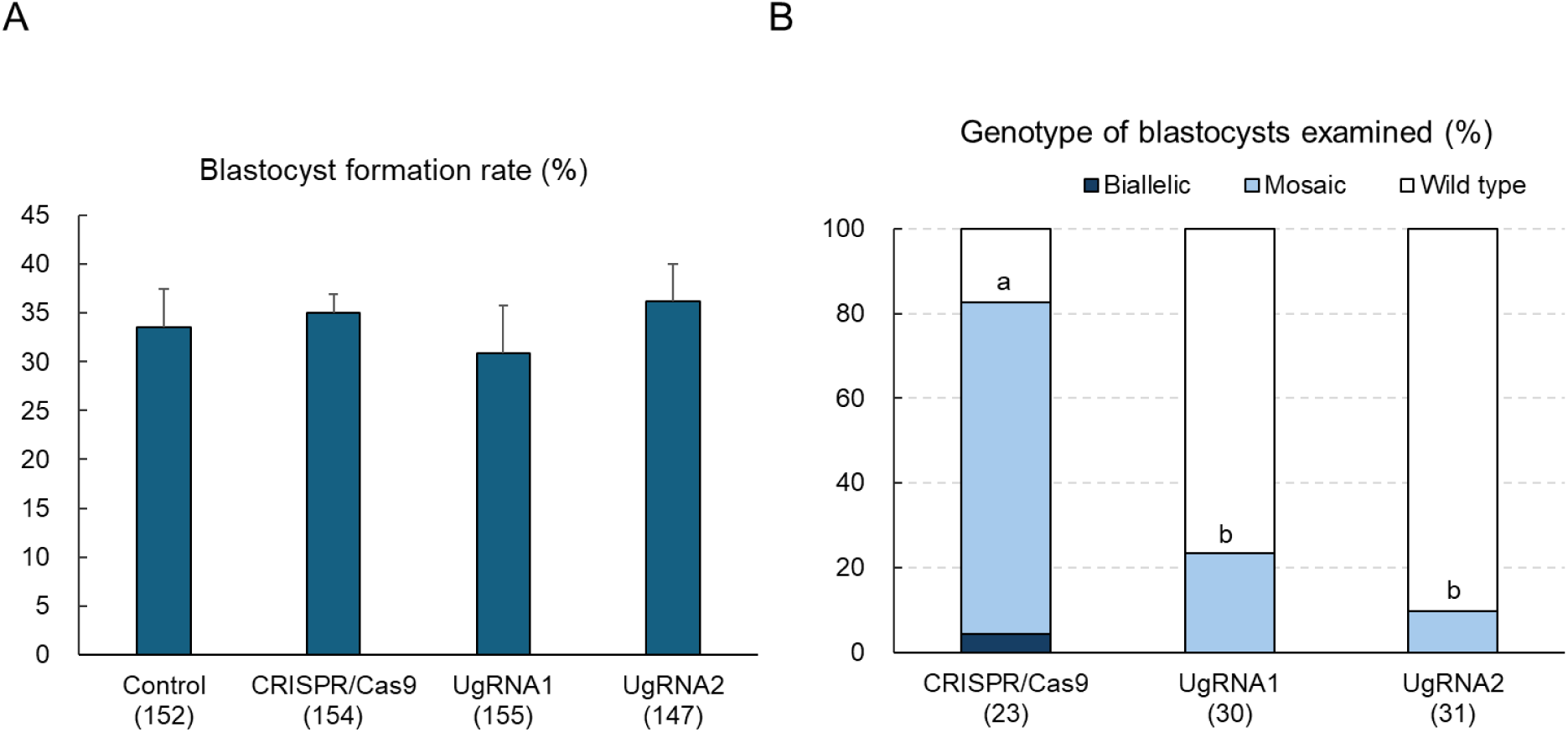
Genome editing via electroporation-mediated introduction of FokI–Nova protein and UgRNA targeting GGTA1. (A) Rate of blastocyst formation in electroporated zygotes. For each treatment group, four replicates of 147–155 oocytes per treatment were analyzed. Numbers in parentheses indicate the total number of treated zygotes. Values of means ± SEM are shown. (B) Blastocyst genotypes after electroporation-mediated introduction of FokI–Nova protein and UgRNA targeting *GGTA1*. The percentage of blastocysts was defined as the ratio of the mutant blastocysts to the total number of blastocysts. For comparison, a control group, in which CRISPR/Cas9 ribonucleoprotein (RNP) complexes were introduced by electroporation, was included. The percentages of mutant blastocysts were analyzed using the chi-square test. ^a–b^ Values with different superscripts differ significantly (*p* < 0.05). WT, wild type; Biallelic, biallelic mutant; Mosaic, mosaic mutant. Numbers in parentheses indicate the total number of blastocysts analyzed.

Therefore, electroporation is a viable alternative to microinjection for delivering UCAY components into porcine zygotes, although mosaicism remains common under current conditions. However, the genome editing efficiency following electroporation was lower than that achieved by microinjection. The efficiency of electroporation-mediated intracellular delivery is affected by several cargo properties, including molecular size, net charge, and concentration [17, 18]. Because the UCAY system designed in this study requires the simultaneous delivery of UgRNA and FokI–Nova fusion proteins, sufficient amounts of both components may not have been introduced into the zygotes under the present conditions. Optimization of the concentrations and molar ratio of the UCAY components, and the electrical pulse parameters, may improve genome editing efficiency. Electroporation did not cause any apparent developmental toxicity. Further optimization of the delivery conditions is required to increase editing efficiency and reduce mosaicism.

## DISCUSSION

In contrast to the CRISPR/Cas9 system, in which the Cas9 protein directly binds to gRNA, the UCAY system uses an indirect interaction mediated by an RNA-binding protein such as Nova. This architecture enables a three-step mechanism for sequence-specific targeting: (i) the guide RNA hybridizes to the target DNA sequence; (ii) an RNA-binding protein recognizes and binds to a distinct region of the guide RNA; and (iii) an effector enzyme fused to the RNA-binding protein acts in proximity to the target site. This modular and sequential process allowed precise access to specific genomic loci.

The UCAY system offers several advantages. First, its simplified licensing framework may facilitate broader commercial applications than CRISPR-based technologies. Secondly, the length of the guide RNA is flexible and not constrained to a fixed size, for example, approximately 20 bp, as in CRISPR/Cas9, allowing for greater adaptability in target design. Third, the UCAY does not require PAM sequences, expanding the range of accessible genomic sites. The system is compatible with repetitive sequences that are often difficult to target using existing technologies. Finally, the relatively small size of protein components may be advantageous for delivery and expression.

Mechanistically, UCAY is a design framework in which target recognition and effector functions are decoupled into separate modules: an RNA component that determines target specificity and a fusion protein that executes the desired molecular activity. The system is not limited to a specific protein or function but instead embodies a modular design principle. In principle, the effector domain of the fusion protein can be replaced by a variety of functional modules, including nucleases, for example, FokI, base-editing enzymes, such as deaminases, epigenetic modifiers, including TET family proteins, and transcriptional regulators that activate or repress gene expression. Similarly, Nova represents one example of an RNA-binding domain, and alternative RNA-binding proteins with different recognition motifs can be used. This flexibility highlights the extensibility of the UCAY platform.

### Limitations of the study

Despite these advantages, this study has several limitations. One major concern is the lack of a full evaluation of off-target effects. Nonspecific cleavage at unintended genomic sites is a critical issue in all genome editing technologies [19, 20]. Previous studies have reported varying degrees of off-target activity in CRISPR/Cas9-edited cells [21, 22] and detectable off-target events in gene-modified mice identified by whole-genome sequencing [23]. However, such events appear to be rare in nonhuman primates [24]. Therefore, a thorough assessment of off-target effects of the UCAY system is essential for its safe and reliable application.

Another limitation is the relatively low editing efficiency observed in porcine zygotes, in which a high degree of mosaicism was detected. Mosaicism is also a known challenge in CRISPR/Cas9-mediated genome editing following zygote microinjection [25, 26] and is thought to arise from the prolonged activity of editing components during early embryonic development [27]. The presence of multiple genotypes within a single organism complicates downstream analyses and necessitates additional breeding. Further optimization is required to improve editing efficiency and reduce mosaicism in the UCAY system. Because UCAY is a newly developed platform, robust design tools have yet to be established. Furthermore, although the length of the target recognition sequence can be flexibly adjusted, further investigation is needed to determine the optimal length for efficient genome editing. Efficient targeting also requires computational methods to identify optimal guide RNA sequences from a large pool of candidates. The development of such design frameworks, incorporating both target recognition sequence length and guide RNA sequence selection, will be critical for maximizing the performance and usability of the UCAY system.

## CONCLUSION

In this study, we established UCAY as an RNA-guided genome-editing platform. Although several limitations remain and require further investigation, this system offers high flexibility in effector function and minimal constraints on target site selection. These features suggest that UCAY has strong potential for a wide range of applications, including future industrial and biomedical applications.

## RESOURCE AVAILABILITY

### Lead contact

Requests for further information and resources should be directed to and will be fulfilled by the lead contact, Fuminori Tanihara and So Makabe.

### Data and code availability

The complete annotated sequences of the plasmids generated in this study are provided in Supplementary Data S1 of GenBank. Any additional information required to reanalyze the data reported in this paper is available from the lead contact upon request.

## ACKNOWLEDGMENTS

We thank Bizan Syokuhin Co. Ltd. Naruto Meat Center (Tokushima, Japan) for supplying pig ovaries. This study was supported in part by KAKENHI grant numbers JP21KK0124, JP23K23805, JP25K09406, and JP25K15936 from the Japan Society for the Promotion of Science (JSPS). The funders played no role in the study design, data collection and analysis, decision to publish, or preparation of the manuscript.

## AUTHOR CONTRIBUTIONS

F.T. and S.M. conceived the study and drafted the manuscript. S.T.N. and S.M. performed most of the experiments. S.M. designed and optimized functional UCAY proteins and gRNAs and validated and analyzed the UCAY system. S.T.N. evaluated the UCAY system in porcine embryos. I.N. conceived the fundamental concept of the UCAY system. Y.M. coordinated the construction of the UCAY system and its evaluation using mammalian cell lines. F.T. designed the study, coordinated experiments using porcine embryos, and reviewed the manuscript. H.K., Y.K., Y.Y., Y.N., M.M., A.N., M.H., and M.N. contributed to laboratory work and statistical analysis. All authors have read and approved the manuscript.

## ETHICS APPROVAL AND CONSENT TO PARTICIPATE

All animal experiments were approved by the Institutional Animal Care and Use Committee of the Tokushima University (approval number: T28-21).

## DECLARATION OF GENERATIVE AI AND AI-ASSISTED TECHNOLOGIES IN THE WRITING PROCESS

During the preparation of this work, the author(s) used no generative AI and AI-Assisted Technologies.

## SUPPLEMENTAL INFORMATION

Supplementary Data. The plasmids used in this study

Supplementary Information. Supplementary Tables including summary of plasmids, oligonucleotides, primers, and UgRNA targeting sequences with target information used in this study

## METHOD DETAILS

The plasmids used in this study are shown in the Supplementary Data. A summary of the plasmids, oligonucleotides, primers, and UgRNA sequences is listed in Supplementary Tables S1 to S3.

### Cloning and expression of the FokI–Nova fusion protein

An expression vector encoding the FokI–Nova fusion protein (Supplementary Data S1A) was introduced into *Escherichia coli* BL21 (DE3) pLysS-competent cells (Merck, Germany). Briefly, 1 µL of plasmid DNA (10 ng/µL) was mixed with 50 µL of competent cells and incubated on ice for 20 min. Heat shock was performed at 42 °C for 1 min, followed by incubation on ice. The transformed cells were recovered in 250 µL of SOC medium and incubated at 37 °C for 1 h. Subsequently, 100 µL of the culture was spread onto LB agar plates containing kanamycin and incubated overnight at 37 °C. Single colonies were selected and cultured overnight at 37 °C. The culture was mixed with an equal volume of 50% glycerol and stored at −80 °C as a glycerol stock.

For protein expression, bacteria from the glycerol stock were inoculated into LB medium containing kanamycin and cultured at 37 °C until the optical density reached approximately 0.6. Protein expression was induced by adding IPTG to a final concentration of 1 mM, followed by incubation at 25 °C overnight. Cells were harvested and resuspended in BugBuster Protein Extraction Reagent (Novagen, Madison, WI, USA) supplemented with 10 µL of Lysonase Bioprocessing Reagent (Novagen) and incubated at 25 °C for 5 min with gentle agitation. The lysate was centrifuged and the supernatant was collected as the soluble fraction.

Purification was performed using the TALON Metal Affinity Resin (Takara Bio, Shiga, Japan). The resin (1 mL) was equilibrated with 10 bed volumes of equilibration buffer (50 mM sodium phosphate and 300 mM sodium chloride, pH 7.4). The soluble fraction was applied to the column to allow binding of the His-tagged fusion protein. The column was washed with eight bed volumes of the equilibration buffer, followed by seven bed volumes of the equilibration buffer containing 10 mM imidazole. The fusion protein was eluted with three bed volumes of elution buffer (50 mM sodium phosphate, 300 mM sodium chloride, and 150 mM imidazole, pH 7.4). Purified protein samples were mixed with EzApply (ATTO, Tokyo, Japan), heat-denatured at 95 °C for 5 min and analyzed using SDS-PAGE on a 5–20% polyacrylamide gel (20 mA, 70 min). Protein expression was confirmed by immunodetection with an anti-FokI antibody.

### Design and preparation of UgRNA targeting GFP

The sequence of UgRNA targeting GFP (UgRNA-GFP) is shown in Supplementary Fig. S1. UgRNAs were constructed as follows:

#### Hybridization region complementary to the target sequence

Using the pEGFP-N1 plasmid (Takara Bio) (Supplementary Data S3) as a template, a 290 bp fragment corresponding to the GFP target region was amplified by PCR using the following primers: eGFP_Xba5P (forward) and eGFP_Spe3P (reverse). Restriction enzyme sites (XbaI and SpeI) were introduced at both ends. The PCR products were ligated into the pCR2.1 vector using a standard cloning procedure.

#### Nova protein-binding guide region

Oligonucleotides UCAY5P and UCAY3P (Supplementary Table S2) were annealed and extended to generate a guide region. The resulting fragment was digested with BamHI and BglII and ligated downstream of the hybridization region in the pCR2.1 vector (Supplementary Data S1B).

#### In vitro transcription

The plasmid was linearized using HindIII and subjected to in vitro transcription using the T7 Transcription Kit. The resulting UgRNA contains two UCAY repeat motifs derived from α2-glyR, connected via linker sequences [7].

### In vitro cleavage assay of GFP

#### Cleavage assay using PCR-amplified GFP fragments

A GFP fragment was amplified from pEGFP-N1 using the primers GFP_assay_F (forward) and GFP_assay_R (reverse). UgRNA-GFP was incubated with a PCR-amplified GFP fragment containing the target site and FokI–Nova protein at 37 °C for 2 h. Cleavage of the PCR fragment was assessed using a microchip electrophoresis system (MCE-202 MultiNA; Shimadzu, Kyoto, Japan) and agarose gel electrophoresis. Following agarose gel electrophoresis, gel regions excluding the band presumed to represent the uncleaved PCR-amplified GFP fragment were excised and DNA was extracted. The recovered DNA fragments were cloned using the Quick Blunting Kit (New England Biolabs) according to the manufacturer’s instructions, and individual clones were sequenced. Sequencing results were analyzed using SnapGene software (Dotmatics, Boston, MA, USA). The resulting PCR product and design of the in vitro cleavage substrate are shown in Supplementary Fig. S2.

#### Genome editing in 293/GFP cells via RNP delivery

The UgRNA-GFP and FokI–Nova proteins were assembled into RNP complexes before electroporation. Briefly, 1 µg of UgRNA-GFP was mixed with Duplex Buffer (Integrated DNA Technologies; IDT, Coralville, IA, USA), heated at 95 °C for 5 min, and then cooled to 25 °C. Subsequently, 0.5 µg of UCAY nuclease was added, the total volume was adjusted to 10 µL, and the mixture was incubated at 25 °C for 5 min to allow RNP complex formation. For electroporation, 1 × 10^6^ 293/GFP cells were suspended in 90 µL of Opti-MEM I medium and mixed with 10 µL of Duplex buffer containing the RNP complexes (1 µg of UgRNA-GFP and 0.5 µg of UCAY nuclease), resulting in a total electroporation volume of 100 µL. Electroporation was performed in decay mode under the following conditions: poration pulse (Pp), 125 V for 5 ms; interval between poration and driving pulses, 50 ms; driving pulse (Pd), 20 V for 50 ms; driving pulse interval, 50 ms; driving pulse cycles, 5; capacitance, 940 µF; and pulse polarity, +/−. Following electroporation, the cells were cultured for approximately 10 d and passaged every 3 d at a 1:10 dilution. The loss of GFP fluorescence was subsequently evaluated using flow cytometry.

### Flow cytometric analysis and cell sorting

Following the electroporation-mediated introduction of UgRNA-GFP and FokI–Nova proteins, 293/GFP cells were subjected to flow cytometry and cell sorting using a BD FACSMelody^TM^ Cell Sorter (BD Biosciences, San Jose, CA, USA). Cells provided for sorting were recovered from frozen stocks and cultured in high-glucose Dulbecco’s modified Eagle’s medium supplemented with 10% fetal bovine serum, 100 U/mL penicillin, and 100 µg/mL streptomycin at 37 °C under 5% CO_2_. The cells were expanded by three passages before sorting. Singlet cells were gated based on forward scatter measurements and GFP-negative and GFP-positive populations were distinguished based on GFP fluorescence intensity. Non-GFP-expressing HEK293T cells were used as a reference to define GFP-negative gates. The sorting gate was refined after preliminary analyses to improve the recovery of GFP-negative cells. The percentage of GFP-negative cells was calculated relative to that of the gated singlet population. After sorting, aliquots of GFP-negative and GFP-positive fractions were examined by fluorescence microscopy using a Leica DMi8 fluorescence microscope (Leica Microsystems, Wetzlar, Germany) to verify the absence or presence of GFP fluorescence.

### Design and preparation of UgRNA targeting porcine GGTA1

UgRNA1 and 2 targeting porcine *GGTA1* (Supplementary Table S3) were designed and prepared using a procedure similar to that described above for the preparation of UgRNAs targeting GFP, including the hybridization and guide regions. To construct the hybridization region, pUC57-kan_T7-BsaI-UCAY gRNA_v0 (Supplementary Data S1C) was used as the cloning vector for UgRNA transcription. Oligonucleotides corresponding to the designed target-hybridizing sequences (Supplementary Table S2) were annealed to generate complementary 4-bp overhangs at both ends, compatible with the BsaI-digested vector ends, and the resulting fragments were cloned into the vector (Supplementary Data S1D, E). The resulting plasmids were linearized with HindIII and used as templates for in vitro transcription.

Synthesis of the guide region and in vitro transcription of UgRNA were performed in the same manner as described in the section “Design and preparation of UgRNA targeting GFP.”

### Oocyte collection, in vitro maturation, and fertilization

Pig ovaries were obtained from prepubertal gilts (Landrace × Large White × Duroc breeds) at a local slaughterhouse and were transported in physiological saline within 1 h to the laboratory at 30 °C. Ovaries were washed three times with prewarmed physiological saline solution supplemented with 100 IU/ml penicillin G potassium (Meiji, Tokyo, Japan) and 0.1 mg/ml streptomycin sulfate (Meiji). Follicles with diameters of 3–6 mm on the ovarian surface were sliced in sterilized dishes using a surgical blade and cumulus-oocyte complexes (COCs) were visualized and collected under a stereomicroscope. Approximately 50 COCs were cultured in 500 µl of maturation medium consisting of tissue culture medium 199 with Earle’s salts (TCM 199; Gibco/Invitrogen Co., Carlsbad, CA, USA) supplemented with 10% (v/v) porcine follicular fluid, 0.6 mM cysteine (Sigma-Aldrich, St. Louis, MO, USA), 50 µM β-mercaptoethanol (Wako Pure Chemical Industries Ltd., Osaka, Japan), 50 µM sodium pyruvate (Sigma-Aldrich), 2 mg/ml d-sorbitol (Wako Pure Chemical Industries Ltd.), 10 IU/ml equine chorionic gonadotropin (eCG; Kyoritu Seiyaku, Tokyo, Japan), 10 IU/ml human chorionic gonadotropin (hCG; Kyoritu Seiyaku), and 50 µg/ml gentamicin (Sigma-Aldrich), then covered with mineral oil (Sigma-Aldrich) for 22 h in 4-well dishes (Nunc A/S, Roskilde, Denmark). The COCs were transferred to a maturation medium without hormones for an additional 22 h. COCs were incubated at 39 °C in a humidified incubator containing 5% CO_2._

Mature oocytes were subjected to in vitro fertilization as previously described [28]. Briefly, frozen-thawed ejaculated spermatozoa were transferred to 5 ml of porcine fertilization medium (PFM; Research Institute for the Functional Peptides Co., Yamagata, Japan) and washed by centrifugation at 500 × *g* for 5 min. The pelleted spermatozoa were resuspended in the fertilization medium and adjusted to a density of 1 × 10^6^ cells/ml. Approximately 50 oocytes were transferred to 500 µl of sperm-containing fertilization medium, covered with mineral oil in 4-well dishes, and co-incubated for 5 h at 39 °C in a humidified incubator containing 5% CO_2,_ 5% O_2_, and 90% N_2_. After co-incubation, the putative zygotes were denuded from the cumulus cells, attached to spermatozoa by mechanical pipetting, transferred to porcine zygote medium (PZM-5; Research Institute for Functional Peptides Co.), and cultured until cytoplasmic microinjection (CI) or electroporation.

### Cytoplasmic microinjection into porcine zygotes

Presumptive zygotes were subjected to CI of the genome-editing components. Microinjection was performed in a 20 μl drop of PZM-5 (Research Institute for the Functional Peptides Co.) covered by mineral oil. Nuclease-free pure water containing 100 ng/µl UgRNA targeting porcine *GGTA1* and 100 ng/µl FokI–Nova protein, nuclease-free pure water containing 10ng/µl all-in-one plasmid coding both UgRNA targeting porcine *GGTA1* and FokI–Nova protein, or duplex buffer containing 100 ng/µl Cas9 protein (Guide-it™ Recombinant Cas9; Takara Bio) and 100 ng/µl gRNA (Alt-R™ CRISPR crRNAs and tracrRNA, chemically modified and length-optimized variants of native guide RNAs purchased from IDT) [29] (The target sequence is shown in Fig. 3A), was used. The injection medium containing the genome editors was loaded into an injection pipette (Femtotips II; Eppendorf, Hamburg, Germany). Zygotes were immobilized using a holding pipette and the tip of the injection pipette was inserted into the cytoplasm through the zona pellucida and cell membrane at the 3 o’clock position. Subsequently, a solution containing genome editors was injected into the cytoplasm under air pressure (injection pressure, 100–200 hectopascals (hPa); compensation pressure, 10–15 hPa; injection time, 0.15 s) using a microinjector (FemtoJet 4i; Eppendorf). We confirmed the success of CI by observing a slight swelling of the oocyte/zygote cytoplasm.

The zygotes were subsequently transferred to PZM-5. Approximately 50 zygotes were cultured continuously in 500 µl of PZM-5 covered with mineral oil for 7 d in 4-well dishes. The incubation of zygotes and embryos was conducted at 39 °C in a humidified incubator containing 5% CO_2,_ 5% O_2_, and 90% N_2_. To evaluate the genome editing efficiency, blastocysts from microinjected zygotes were collected and subjected to gene analysis, as described below.

### Electroporation

Electroporation was performed as previously described [16]. In brief, an electrode (LF501PT1-20; BEX) was connected to a CUY21EDIT II electroporator (BEX) and placed under a stereoscopic microscope. At 10 h after the start of IVF, the 50 inseminated zygotes were washed with Opti-MEM I solution (Gibco/Invitrogen, Carlsbad, CA, USA) and were aligned within the electrode gap of a chamber slide filled with 10 μl of Nuclease-Free Duplex Buffer (IDT) containing 100 ng/µl UgRNA targeting porcine *GGTA1* (Fig. 3A and Supplementary Fig. S1) and 100 ng/µl FokI–Nova protein or 100 ng/μl gRNA (Alt-R CRISPR crRNAs and tracrRNA) targeting *GGTA1* [29] (The target sequence is shown in Figure 3A) and 100 ng/μl Cas9 protein (Takara Bio). After electroporation (five 1-ms square pulses at 25 V), the zygotes were washed and cultured using PZM-5, as described above. To evaluate the genome editing efficiency, blastocysts from electroporated zygotes were collected and subjected to gene analysis, as described below.

### Analysis of targeted genes in porcine blastocysts

Genomic DNA was isolated from blastocysts by boiling in a 50 mM NaOH solution. After neutralization, the genomic regions flanking the gRNA target sequences were PCR-amplified using specific primers targeting exon 1 of the *GGTA1* gene, including the target site of the gRNA: GGTA1_genotype F (forward) and GGTA1_genotype R (reverse). PCR products from blastocysts were extracted using agarose gel electrophoresis. The target genomic regions of the PCR products were sequenced directly. Sanger sequencing was performed using a BigDye Terminator Cycle Sequencing Kit ver. 3.1 (Thermo Fisher Scientific, Waltham, MA, USA) and an ABI 3500 Genetic Analyzer (Applied Biosystems, Foster City, CA, USA). The TIDE bioinformatics package [15] was used to determine genome editing efficiency in blastocysts. Blastocysts carrying no WT sequences were classified as having biallelic mutations. Blastocysts carrying one or more mutations in the WT sequence were classified as mosaics. Blastocysts carrying only the WT sequence were classified as WT. The mutation efficiency was defined as the ratio of the number of blastocysts carrying mutations to the total number of blastocysts examined.

### Quantification and statistical analysis

Percentage data for embryos that developed to the blastocyst stage were subjected to arcsine transformation before analysis of variance (ANOVA). The transformed data were evaluated using ANOVA, followed by the protected Fisher’s least significant difference test. The analysis was performed using the StatView software (Abacus Concepts, Berkeley, CA, USA). The percentage of mutated blastocysts was analyzed using the chi-square test. Differences with a probability value (*p*) of 0.05 or less were considered statistically significant.

---

**Supplementary Table S1.**
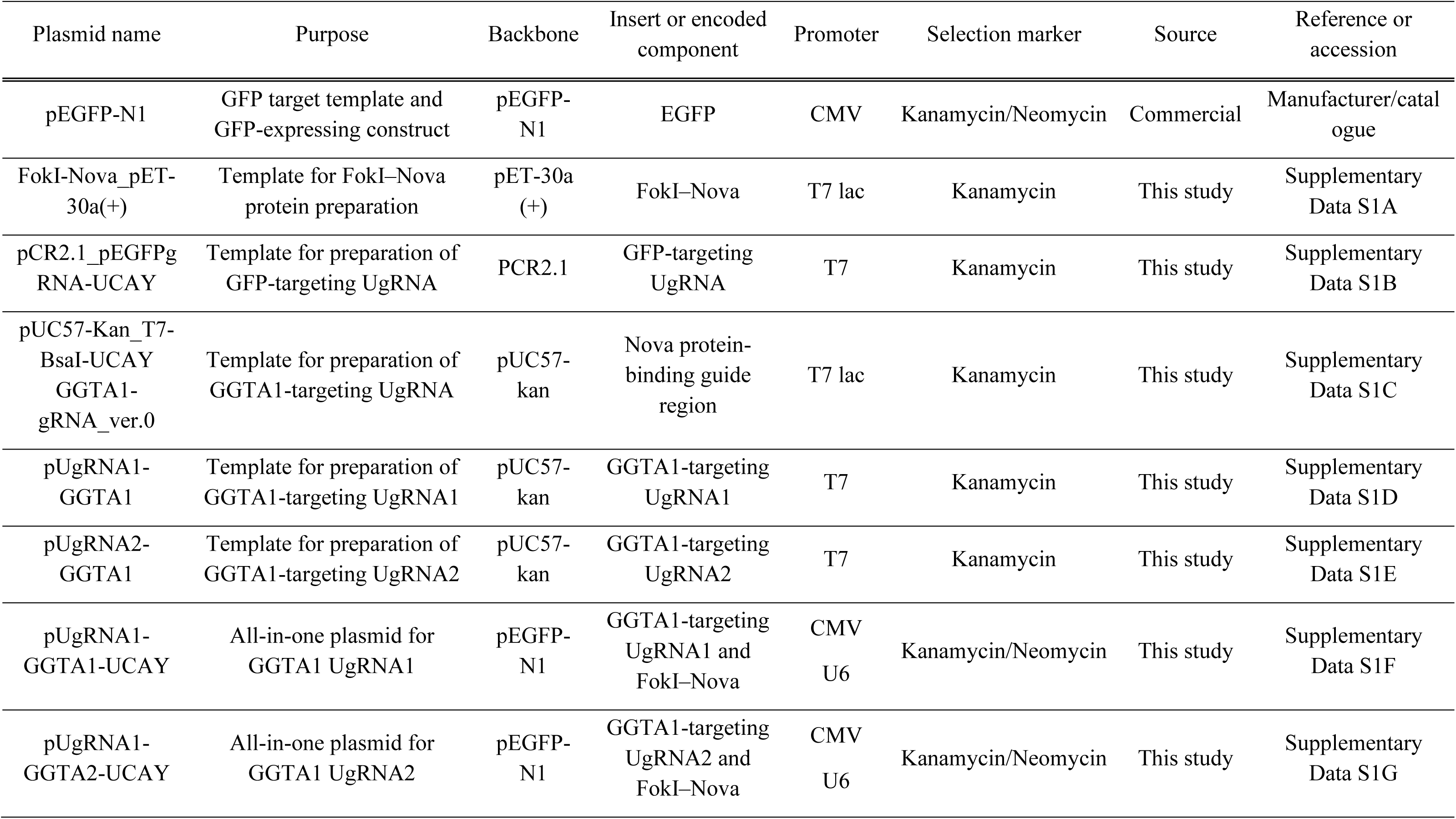
Plasmids used in this study.

**Supplementary Table S2.**
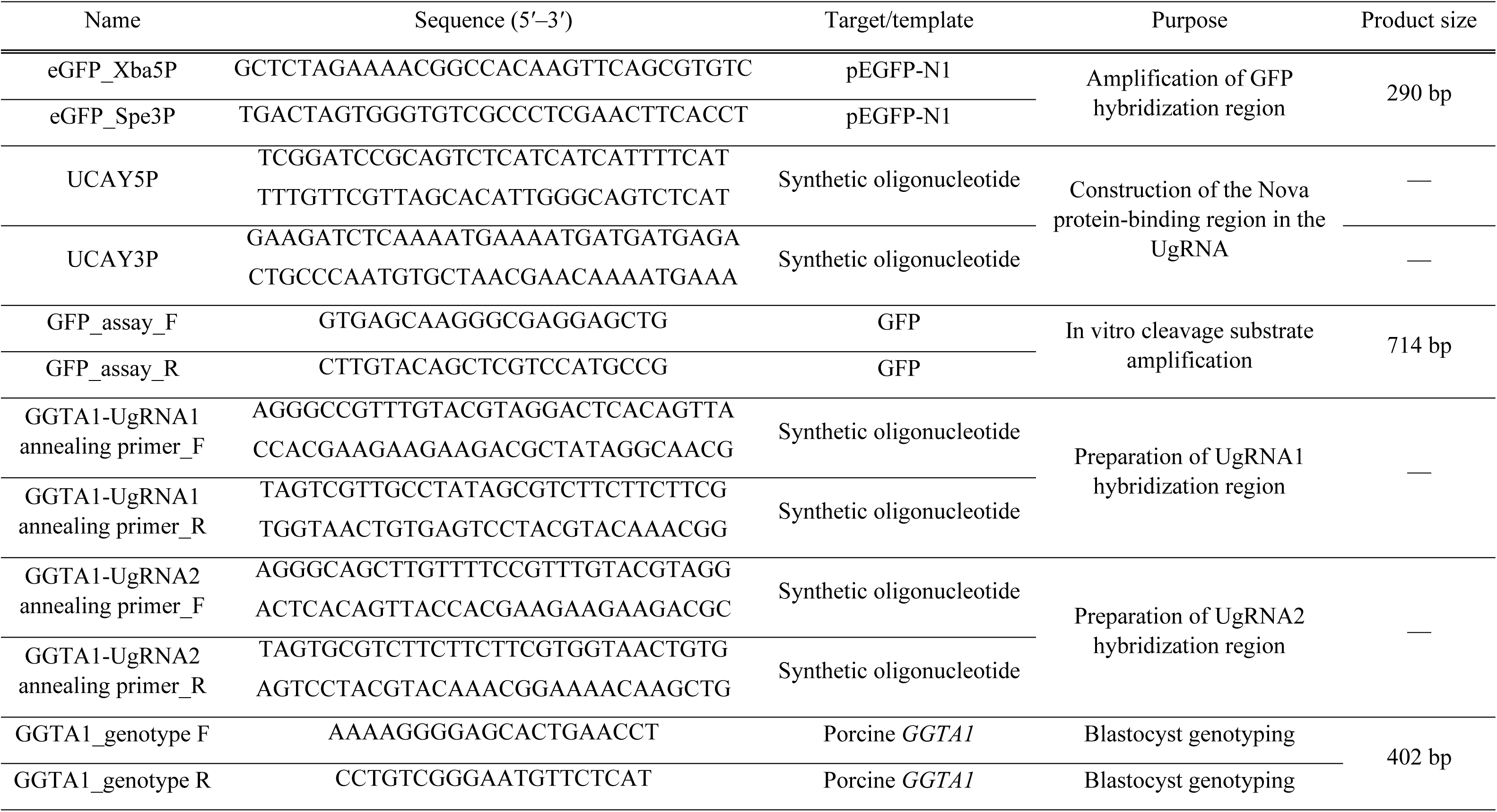
Oligonucleotides and primers used in this study.

**Supplementary Table S3.**
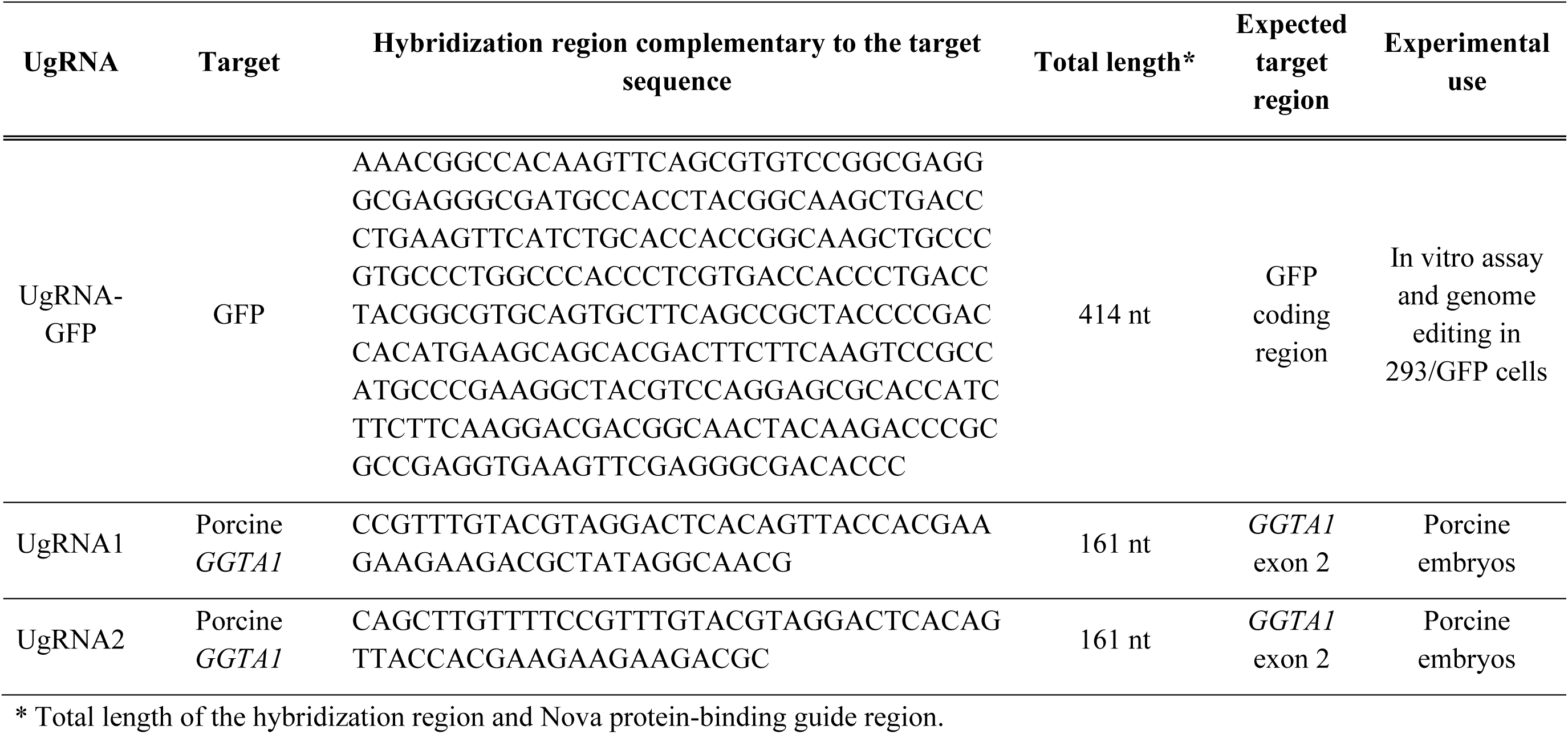
UgRNA targeting sequences and target information.

## Supplementary Figures

**Supplementary Figure S1.**
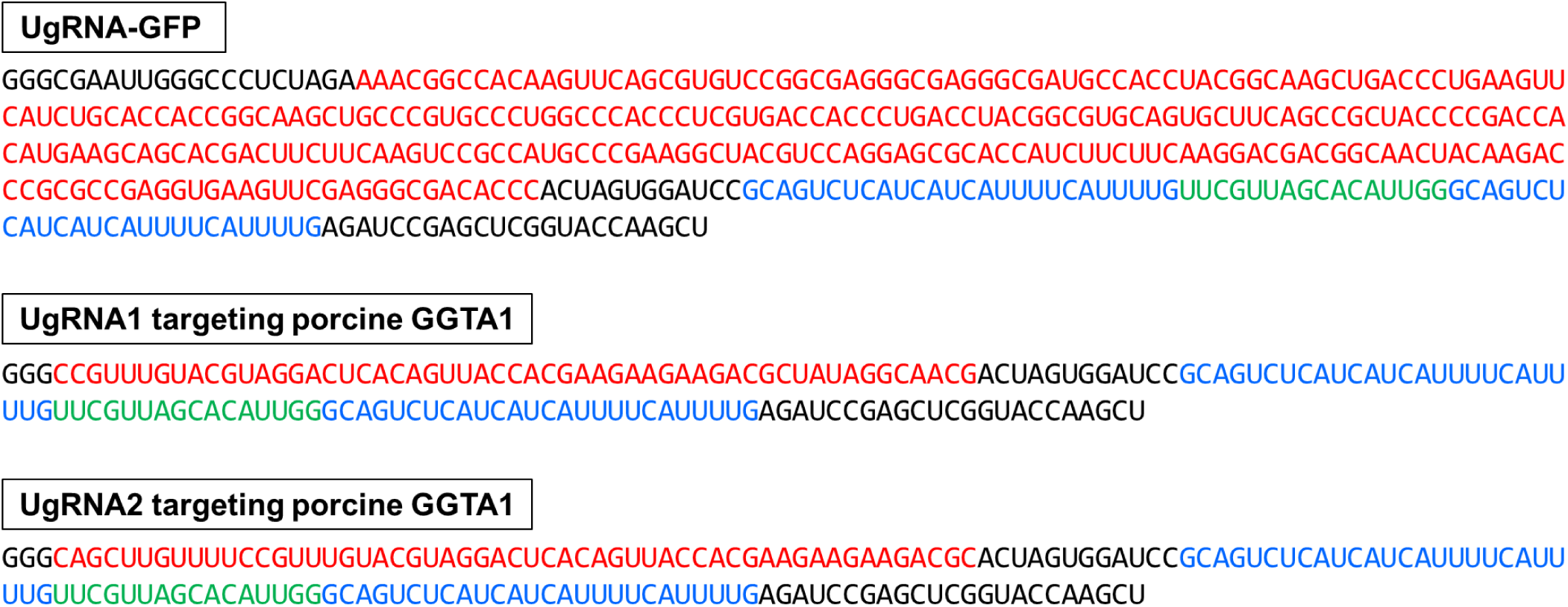
UgRNA sequences used in this study. Red text indicates a sequence complementary to the target, blue text indicates the UCAY repeat sequence, and green text indicates the UCAY repeat spacer sequence.

**Supplementary Figure S2.**
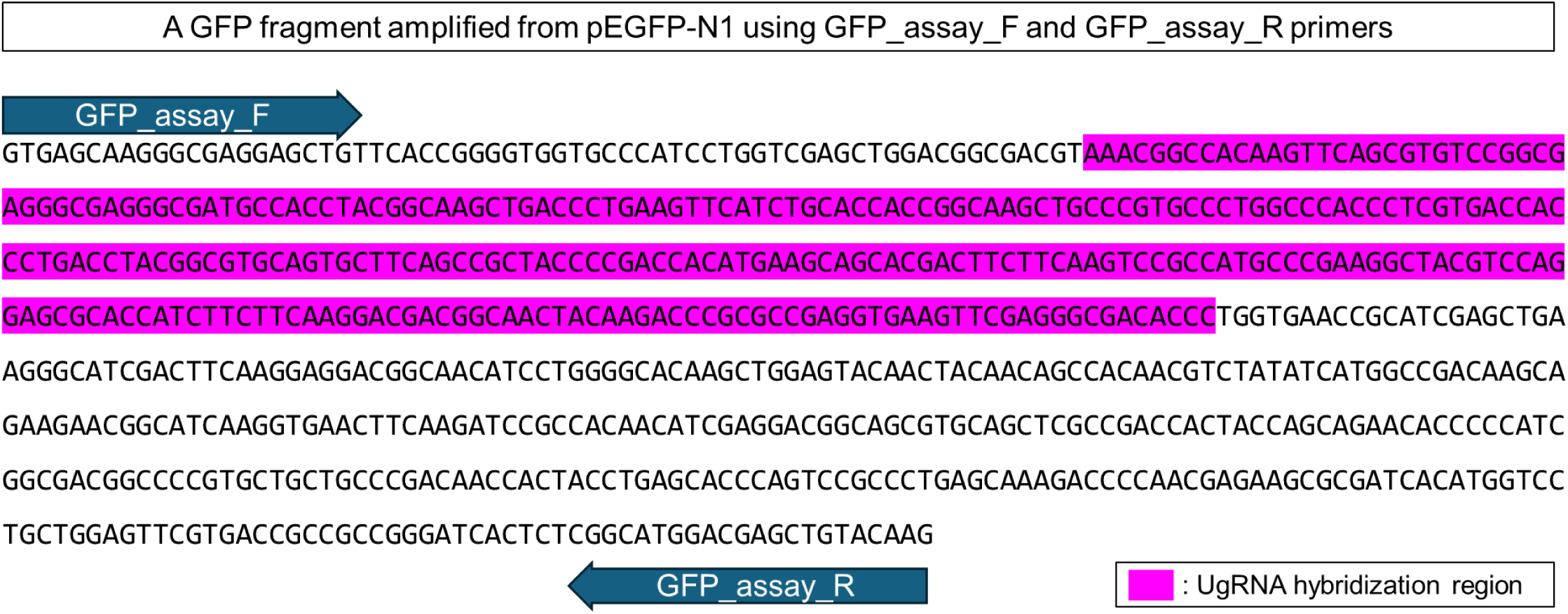
GFP target sequence and in vitro cleavage-substrate design.

## Supplementary Data

**Supplementary Data S1. Annotated complete sequence of the FokI–Nova and UgRNA expression plasmid**

Maps are in genbank format in a ZIP file. (A) FokI–Nova fusion protein expression plasmid (FokI-Nova_pET-30a(+)). (B) Plasmid used for in vitro transcription of UgRNA targeting GFP. (C) Plasmid used for in vitro transcription of UgRNA targeting GGTA1 (pUC57-Kan_T7-BsaI-UCAY GGTA1-gRNA_ver.0). (D) Plasmid used for in vitro transcription of UgRNA1 targeting GGTA1 (pUgRNA1-GGTA1). (E) Plasmid used for in vitro transcription of UgRNA2 targeting GGTA1 (pUgRNA2-GGTA1). (F) FokI–Nova and UgRNA1 targeting GGTA1 expression all-in-one plasmid (pUgRNA1-GGTA1-UCAY). (G) FokI–Nova and UgRNA2 targeting GGTA1 expression all-in-one plasmid (pUgRNA2-GGTA1-UCAY).

**Supplementaly Data S2. Amino acid sequence of FokI-Nova fusion protein**

MHHHHHHSSGLVPRGSGMKETAAAKFERQHMDSPDLGTDDDDKAMADIGSQLVKSEL EEKKSELRHKLKYVPHEYIELIEIARNSTQDRILEMKVMEFFMKVYGYRGKHLGGSRKP DGAIYTVGSPIDYGVIVDTKAYSGGYNLPIGQADEMQRYVEENQTRNKHINPNEWWKV YPSSVTEFKFLFVSGHFKGNYKAQLTRLNHITNCNGAVLSVEELLIGGEMIKAGTLTLEE VRRKFNNGEINFSGSETPGTSESATPESKDVVEIAVPENLVGAILGKGGKTLVEYQELTG ARIQISKKGEFVPGTRNRKVTITGTPAATQAAQYLITQRITYEQGVRAANPQKVGSRADP KKKRKV*

**Supplementaly Data S3. The sequence of pEGFP-N1 plasmid**

TAGTTATTAATAGTAATCAATTACGGGGTCATTAGTTCATAGCCCATATATGGAGTT CCGCGTTACATAACTTACGGTAAATGGCCCGCCTGGCTGACCGCCCAACGACCCCCG CCCATTGACGTCAATAATGACGTATGTTCCCATAGTAACGCCAATAGGGACTTTCCA TTGACGTCAATGGGTGGAGTATTTACGGTAAACTGCCCACTTGGCAGTACATCAAGT GTATCATATGCCAAGTACGCCCCCTATTGACGTCAATGACGGTAAATGGCCCGCCTG GCATTATGCCCAGTACATGACCTTATGGGACTTTCCTACTTGGCAGTACATCTACGT ATTAGTCATCGCTATTACCATGGTGATGCGGTTTTGGCAGTACATCAATGGGCGTGG ATAGCGGTTTGACTCACGGGGATTTCCAAGTCTCCACCCCATTGACGTCAATGGGAG TTTGTTTTGGCACCAAAATCAACGGGACTTTCCAAAATGTCGTAACAACTCCGCCCC ATTGACGCAAATGGGCGGTAGGCGTGTACGGTGGGAGGTCTATATAAGCAGAGCTG GTTTAGTGAACCGTCAGATCCGCTAGCGCTACCGGACTCAGATCTCGAGCTCAAGCT TCGAATTCTGCAGTCGACGGTACCGCGGGCCCGGGATCCACCGGTCGCCACCATGGT GAGCAAGGGCGAGGAGCTGTTCACCGGGGTGGTGCCCATCCTGGTCGAGCTGGACG GCGACGTAAACGGCCACAAGTTCAGCGTGTCCGGCGAGGGCGAGGGCGATGCCACC TACGGCAAGCTGACCCTGAAGTTCATCTGCACCACCGGCAAGCTGCCCGTGCCCTGG CCCACCCTCGTGACCACCCTGACCTACGGCGTGCAGTGCTTCAGCCGCTACCCCGAC CACATGAAGCAGCACGACTTCTTCAAGTCCGCCATGCCCGAAGGCTACGTCCAGGA GCGCACCATCTTCTTCAAGGACGACGGCAACTACAAGACCCGCGCCGAGGTGAAGT TCGAGGGCGACACCCTGGTGAACCGCATCGAGCTGAAGGGCATCGACTTCAAGGAG GACGGCAACATCCTGGGGCACAAGCTGGAGTACAACTACAACAGCCACAACGTCTA TATCATGGCCGACAAGCAGAAGAACGGCATCAAGGTGAACTTCAAGATCCGCCACA ACATCGAGGACGGCAGCGTGCAGCTCGCCGACCACTACCAGCAGAACACCCCCATC GGCGACGGCCCCGTGCTGCTGCCCGACAACCACTACCTGAGCACCCAGTCCGCCCTG AGCAAAGACCCCAACGAGAAGCGCGATCACATGGTCCTGCTGGAGTTCGTGACCGC CGCCGGGATCACTCTCGGCATGGACGAGCTGTACAAGTAAAGCGGCCGCGACTCTA GATCATAATCAGCCATACCACATTTGTAGAGGTTTTACTTGCTTTAAAAAACCTCCC ACACCTCCCCCTGAACCTGAAACATAAAATGAATGCAATTGTTGTTGTTAACTTGTT TATTGCAGCTTATAATGGTTACAAATAAAGCAATAGCATCACAAATTTCACAAATAA AGCATTTTTTTCACTGCATTCTAGTTGTGGTTTGTCCAAACTCATCAATGTATCTTAA GGCGTAAATTGTAAGCGTTAATATTTTGTTAAAATTCGCGTTAAATTTTTGTTAAATC AGCTCATTTTTTAACCAATAGGCCGAAATCGGCAAAATCCCTTATAAATCAAAAGAA TAGACCGAGATAGGGTTGAGTGTTGTTCCAGTTTGGAACAAGAGTCCACTATTAAAG AACGTGGACTCCAACGTCAAAGGGCGAAAAACCGTCTATCAGGGCGATGGCCCACT ACGTGAACCATCACCCTAATCAAGTTTTTTGGGGTCGAGGTGCCGTAAAGCACTAAA TCGGAACCCTAAAGGGAGCCCCCGATTTAGAGCTTGACGGGGAAAGCCGGCGAACG TGGCGAGAAAGGAAGGGAAGAAAGCGAAAGGAGCGGGCGCTAGGGCGCTGGCAAG TGTAGCGGTCACGCTGCGCGTAACCACCACACCCGCCGCGCTTAATGCGCCGCTACA GGGCGCGTCAGGTGGCACTTTTCGGGGAAATGTGCGCGGAACCCCTATTTGTTTATT TTTCTAAATACATTCAAATATGTATCCGCTCATGAGACAATAACCCTGATAAATGCT TCAATAATATTGAAAAAGGAAGAGTCCTGAGGCGGAAAGAACCAGCTGTGGAATGT GTGTCAGTTAGGGTGTGGAAAGTCCCCAGGCTCCCCAGCAGGCAGAAGTATGCAAA GCATGCATCTCAATTAGTCAGCAACCAGGTGTGGAAAGTCCCCAGGCTCCCCAGCA GGCAGAAGTATGCAAAGCATGCATCTCAATTAGTCAGCAACCATAGTCCCGCCCCT AACTCCGCCCATCCCGCCCCTAACTCCGCCCAGTTCCGCCCATTCTCCGCCCCATGG CTGACTAATTTTTTTTATTTATGCAGAGGCCGAGGCCGCCTCGGCCTCTGAGCTATTC CAGAAGTAGTGAGGAGGCTTTTTTGGAGGCCTAGGCTTTTGCAAAGATCGATCAAG AGACAGGATGAGGATCGTTTCGCATGATTGAACAAGATGGATTGCACGCAGGTTCT CCGGCCGCTTGGGTGGAGAGGCTATTCGGCTATGACTGGGCACAACAGACAATCGG CTGCTCTGATGCCGCCGTGTTCCGGCTGTCAGCGCAGGGGCGCCCGGTTCTTTTTGTC AAGACCGACCTGTCCGGTGCCCTGAATGAACTGCAAGACGAGGCAGCGCGGCTATC GTGGCTGGCCACGACGGGCGTTCCTTGCGCAGCTGTGCTCGACGTTGTCACTGAAGC GGGAAGGGACTGGCTGCTATTGGGCGAAGTGCCGGGGCAGGATCTCCTGTCATCTC ACCTTGCTCCTGCCGAGAAAGTATCCATCATGGCTGATGCAATGCGGCGGCTGCATA CGCTTGATCCGGCTACCTGCCCATTCGACCACCAAGCGAAACATCGCATCGAGCGA GCACGTACTCGGATGGAAGCCGGTCTTGTCGATCAGGATGATCTGGACGAAGAGCA TCAGGGGCTCGCGCCAGCCGAACTGTTCGCCAGGCTCAAGGCGAGCATGCCCGACG GCGAGGATCTCGTCGTGACCCATGGCGATGCCTGCTTGCCGAATATCATGGTGGAAA ATGGCCGCTTTTCTGGATTCATCGACTGTGGCCGGCTGGGTGTGGCGGACCGCTATC AGGACATAGCGTTGGCTACCCGTGATATTGCTGAAGAGCTTGGCGGCGAATGGGCT GACCGCTTCCTCGTGCTTTACGGTATCGCCGCTCCCGATTCGCAGCGCATCGCCTTCT ATCGCCTTCTTGACGAGTTCTTCTGAGCGGGACTCTGGGGTTCGAAATGACCGACCA AGCGACGCCCAACCTGCCATCACGAGATTTCGATTCCACCGCCGCCTTCTATGAAAG GTTGGGCTTCGGAATCGTTTTCCGGGACGCCGGCTGGATGATCCTCCAGCGCGGGGA TCTCATGCTGGAGTTCTTCGCCCACCCTAGGGGGAGGCTAACTGAAACACGGAAGG AGACAATACCGGAAGGAACCCGCGCTATGACGGCAATAAAAAGACAGAATAAAAC GCACGGTGTTGGGTCGTTTGTTCATAAACGCGGGGTTCGGTCCCAGGGCTGGCACTC TGTCGATACCCCACCGAGACCCCATTGGGGCCAATACGCCCGCGTTTCTTCCTTTTCC CCACCCCACCCCCCAAGTTCGGGTGAAGGCCCAGGGCTCGCAGCCAACGTCGGGGC GGCAGGCCCTGCCATAGCCTCAGGTTACTCATATATACTTTAGATTGATTTAAAACT TCATTTTTAATTTAAAAGGATCTAGGTGAAGATCCTTTTTGATAATCTCATGACCAAA ATCCCTTAACGTGAGTTTTCGTTCCACTGAGCGTCAGACCCCGTAGAAAAGATCAAA GGATCTTCTTGAGATCCTTTTTTTCTGCGCGTAATCTGCTGCTTGCAAACAAAAAAAC CACCGCTACCAGCGGTGGTTTGTTTGCCGGATCAAGAGCTACCAACTCTTTTTCCGA AGGTAACTGGCTTCAGCAGAGCGCAGATACCAAATACTGTCCTTCTAGTGTAGCCGT AGTTAGGCCACCACTTCAAGAACTCTGTAGCACCGCCTACATACCTCGCTCTGCTAA TCCTGTTACCAGTGGCTGCTGCCAGTGGCGATAAGTCGTGTCTTACCGGGTTGGACT CAAGACGATAGTTACCGGATAAGGCGCAGCGGTCGGGCTGAACGGGGGGTTCGTGC ACACAGCCCAGCTTGGAGCGAACGACCTACACCGAACTGAGATACCTACAGCGTGA GCTATGAGAAAGCGCCACGCTTCCCGAAGGGAGAAAGGCGGACAGGTATCCGGTAA GCGGCAGGGTCGGAACAGGAGAGCGCACGAGGGAGCTTCCAGGGGGAAACGCCTG GTATCTTTATAGTCCTGTCGGGTTTCGCCACCTCTGACTTGAGCGTCGATTTTTGTGA TGCTCGTCAGGGGGGCGGAGCCTATGGAAAAACGCCAGCAACGCGGCCTTTTTACG GTTCCTGGCCTTTTGCTGGCCTTTTGCTCACATGTTCTTTCCTGCGTTATCCCCTGATT CTGTGGATAACCGTATTACCGCCATGCAT

